# Effect of *Septoria* brown spot on soybean yield in Illinois

**DOI:** 10.1101/620112

**Authors:** Heng-An Lin, María B. Villamil, Santiago X. Mideros

## Abstract

Brown spot caused by *Septoria glycines* is a prevalent foliar disease in all soybean production areas. Application of foliar fungicides after bloom reduces the disease severity, yet yield responses are not consistent among locations and years. Our research goal was to determine the effect of different levels of *Septoria* brown spot on yield. Different levels of disease severity were effectively obtained in the field by weekly application of chlorothalonil for three, six, and nine times after disease inoculation at V3/V4 stage. Fungicide treatments had a significant effect on vertical progress and chlorotic area with no statistically significant effect on yield. Soybean yield was negatively correlated with vertical progress of the disease (r = −0.36). The vertical progress was the best linear predictor of yield. Based on this model, when the vertical progress of brown spot at R6 increased by 10%, the yield decreased by 142.13 kg/ha (3.4%). A variance component analyses of our data showed that location was the most critical factor, illustrating the significant effect of local environmental conditions on the disease. Power analyses indicated that at least eight locations are needed to detect an effect of 269 kg/ha. Our results provide useful information to improve the experimental design for future experiments addressing the yield constrain by late season diseases of soybean.

## 1 Introduction

Soybean (*Glycine max* (L.) Merr.) is one of the most important crops in the United States with a cultivated area of 35.75 million hectares, and an average yield of 3.5 metric tons/hectare (52.1 bushels/acre) in 2018 [1]. An estimated 12% of soybean yield is lost to diseases every year in the U.S. [2]. *Septoria* brown spot (*Septoria glycines* Hemmi) is one of the ten most destructive diseases associated with the yield losses [3]. This disease often occurs simultaneously with other late-season diseases such as frogeye leaf spot (*Cercospora sojina* Hara), and *Cercospora* leaf blight (*Cercospora kikuchii* (Mat and Tom) Gardner). Some authors consider them a complex of diseases [4]. These late-season diseases have not received much research attention yet assessing their individual effect on soybean yield is critical to precision management practices and our understanding of disease progression in the field.

In this study, we focus on *Septoria* brown spot of soybeans, a highly prevalent disease in all soybean growing areas [2]. This disease was first reported in the US in 1922 [5] and can be found in other top soybean producing countries such as Argentina, Brazil, and China [2]. The typical symptoms are irregular, dark-brown necrotic spots with a yellow halo, that can be observed as early as V2 stage (two fully developed trifoliate leaves on the plant) [6, 7]. Under humid and warm environments, the disease can gradually develop to the upper canopy, leading to premature defoliation and yield losses [8, 9].

Several studies have evaluated the yield losses caused by *Septoria* brown spot. In a field survey with 1000 soybean plant introductions, Lim (1979) [10] showed that yield losses due to *Septoria* brown spot ranged between 1% to 27%, depending on variety, location, and whether the plots were inoculated or not. Yield reductions ranging from 12% to 34% were reported in inoculated plots, and from 8% to 8.7% in naturally infected plots [11]. A study [12] in Ohio reported that yield losses ranged from 2.5% and 9.5% in naturally infected fields with different number of chlorothalonil applications.

No source of resistance to this pathogen has been identified from a survey of 1000 and 626 soybean lines [10, 13]. There is no data on the resistance or susceptibility of newly released soybean cultivars to brown spot [12], and no pathogenic variability among isolates has been found [14].

Application of chlorothalonil (FRAC CODE M05) or mixtures of demethylation inhibitor (FRAC CODE 3) and quinone outside inhibitor (FRAC CODE 11) compounds between R1 to R6 can effectively control *Septoria* brown spot [12]. Fungicide treatments can significantly reduce disease severity at the end of the season; however, the reported impacts on yield varied among locations and years in several studies. Pataky and Lim (1980) [15] reported that spraying benomyl (FRAC CODE 1) at R3 or R3 plus R6 stages had the best effect on reducing disease severity, but the yield increases due to the fungicide application was detected only in one year. Cooper (1989) [16] detected yield increases only among two out of four cultivars when sprayed benomyl, in 2-weeks intervals, between R2 to R6 stages of soybean. Cruz et al. (2010) [12] detected a statistically significant effect of application of strobilurins (pyraclostrobin or azoxysrtobin (FRAC CODE 11)) or a combination of strobilutins and triazole (tebuconazole (FRAC CODE 3)) on reducing disease severity but the yield increases were found only in three out of six location-years.

The amount of photosynthate allocated to the seeds during maturity is the critical factor that determines the seed number in soybeans [17]. Green leaf area duration during this peroid is important for the plant to produce sufficient photosynthate. Therefore, disease control during this peroid is critical for the final yield. Carmona et al. (2011) [4] reported a significant correlation between rain and yield response to fungicide application when rain occurred after pod formation (R3) and before seed formation (R5). These findings suggest that reducing foliar diseases, in general, could have a positive effect on yield under certain environmental circumstances.

While general rules for fungicide application are useful, more detailed knowledge on microorganism-specific yield losses is necessary for the development of precision disease management strategies. The ubiquity of *Septoria* brown spot symptoms suggests that it is a major component of the late-season diseases complex [4]. To accurately calculate the effect of plant pathogens on yield reductions, the priority is to develop reliable methods to exert different levels of disease pressure on field trials. These experiments are often complicated by the large effect of environmental factors on disease development, thus requiring multiple diverse environments to reach valid conclusions. On the other hand, these experiments are expensive and time consuming thus it’s important to know the level of replication necessary to obtain useful results. Our goals were to i) develop a method to obtain different levels of *Septoria* brown spot disease severity on inoculated field trials; ii) determine the effects of *Septoria* brown spot disease severity on soybean yield; and iii) determine the minimum number of replications required for field trial evaluations of the effect of *Septoria* brown spot on crop yields.

## 2 Materials and methods

### 2.1 Field management and experimental layout

Field trials were established at the Crop Sciences Research and Education Center near Urbana (N40.072170°, W88.220882°), the Orr Agricultural Research and Demonstration Center near Perry (N39.805006 °, W90.823208 °), and the Northwestern Illinois Agricultural Research and Demonstration Center near Monmouth, Illinois (N40.931887°, W90.725123°) in 2017. At each location, the fungicide treatment was arranged in a randomized complete block design (RCBD) with four replications. The fungicide treatment had five levels: two controls without fungicide application [control: disease inoculated plots; and NIC: not-inoculated control plots]; and three levels of fungicide 3X, 6X, and 9X, that represent the number of weekly applications performed in order to obtain different levels of disease in the field. Chlorothalonil (Echo^®^ 720 AG, Sipcam Agro USA, Inc.) was applied weekly at a rate of 1.05 kg ai/ha, starting one week after disease inoculation and continuing for 3, 6, or 9 weeks in each of the 3X, 6X, and 9X treatments, respectively. Fungicide was applied using a CO_2_-pressurized backpack sprayer and a 0.48 m 601B-SST - four nozzle light weight boom (R&D Sprayers Bellspray, Inc, USA) with four TJ60-11008 (50) nozzles (TeeJet Technology, USA). Each plot consisted of four 5.2 m long rows, and each row was 0.76 m wide. All trials were planted with soybean cultivar Williams between May 5 and May 18 at 348,480 seeds/hectare. The field in Urbana received supplemental irrigation using a 14VH 1/2” inlet full circle brass wedge drive impact sprinklers (Rain Bird, USA) placed at 1.5 m height, and spaced every 9.14 m.

To prepare the inoculum, three isolates of *Septoria glycines* (R3126, 16S006 and 16S012) were cultured on 3% malt extract agar (MEA) for two weeks at room temperature (~24 °C), with 12hr of fluorescent light. Sterile water was added to each plate, and 10 to 20 μl of the spore suspension was transferred and spread on 3% MEA plates to generate a large quantity of spores. One week later, 5 to 10 ml water amended with 10 μg/L chloramphenicol was added to each plate, and three milliliters was then transferred to a 1 L flask containing 135 g barley and 15 g rice grain. The grain had been prepared by washing 135 g barley and 15 g rice grain with tap water three times and then soaking in the water overnight at 4°C. Excess water was drained before autoclaving. The water content of the substrate prepared in this way, was approximately 60%. Following two weeks of incubation, the conidia were collected with water containing 0.05% Tween 20 and filtered through sterilized, double-layered cheesecloth. The concentration of the spore suspension was adjusted to 10^6^ spores/ml using a hemocytometer. The center two rows of each plot (except the NIC) were inoculated between the V3 and V4 developmental stages [6] with a suprema bak-pak sprayer (Hudson Manufacturing Company, USA).

### 2.2 Disease and yield assessment

Visual ratings (expressed as %) of four components of disease severity (vertical progress, defoliation rate, chlorotic, and necrotic area) were taken weekly from the center two rows of each plot. Vertical progress and defoliation rate were estimated by the percent height reached by brown spot symptoms, or removal of the leaves. The chlorotic and necrotic areas were the visual estimates of the percent of the whole plant foliage that was yellow or black, respectively. To evaluate soybean yield components, twenty plants were randomly collected from each plot. After drying the plants at room temperature, the weight of 100-seeds, the number of pods per plant, and the number of seeds per plant were recorded. Field plots were harvested with a research plot combine between Oct 3 and Oct 18. Soybean yield data (kg/ha) was adjusted to 13% water content. Yield response was calculated by subtracting the yield from treatments (3X, 6X or 9X) to the control plots (0X) in each block.

### 2.3 Data analysis

The area under the disease progress curve (AUDPC) values for the components of disease severity were calculated using the AUDPC function in the Agricolae package [18] in the software R version 3.5.2 [19]. A simple and a multi-location RCBD models were used to evaluate the experiment.

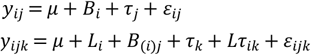

In the simple RCBD model, *y_ij_* is the observed data (yield components and AUDPC for each of the disease severity components) corresponding to the *i^th^* block and *j^th^* treatment. *μ* is the grand population mean, *B_i_* is the random block effect, *τ_j_* is the fixed treatment effect, and *ε_ij_* is the random error term 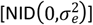. In the multi-location RCBD model, location (L_i_) is added to the model as a random effect with block (B_(i)j_) nested within it. *Lτ_ik_* is the random interaction effect between the i^th^ location and the k^th^ treatment.

Linear mixed models to evaluate the effects of fungicide treatments on yield and disease severity (raw data or AUDPC) were fitted using PROC MIXED in SAS v9.4 [20]. The covtest option was specified in the PROC MIXED statement to estimate the variance components. Least square means were separated using the PDIFF option of LSMEANS in SAS PROC MIXED setting the probability of Type I error or α level (α) = 0.05. Correlation among variables were evaluated with PROC CORR. A stepwise regression analysis (using the REG procedure) was conducted with the components of disease severity as dependent variables and yield as the response variables. Figures were generated in GraphPad Prism Version 6.0 for Windows [21].

### 2.4 Statistical power analysis

Based on our results, we conducted a power analysis using the SIMR package [22] in R [19]. Our goal was to determine the proper number of locations and replicates needed to detect a significant yield response. SIMR produces Monte Carlo stochastic simulations with our empirical data in the linear mixed model indicated above and predetermined effect sizes. For this analysis, we used only the data from the control plots (0X) and the 3X treatment. The threshold of desired power was set as 0.8. The predetermined effect sizes used for the simulations with the yield data were 134.5, 269, and 403.5 kg/ha (two, four, and six bu/ac) which cover the range of 5-10% yield reduction for soybean cultivar Williams. The range of 5 −10% of yield reduction is also the range of reported yield responses to fungicide without disease [23, 24], and with *Septoria* Brown spot [12]. For the other yield components, the effects used for the simulations were 1.5, two, and four pods per plant; three, six, and nine seeds per plant and 0.2, 0.4, and 0.6 grams of 100-seed weight. These values were based on a previous report with *Septoria* brown spot infected assays [9].

## 3 Results

### 3.1 Disease response to fungicide treatments

Weekly evaluations produced nine observations from Perry, and eleven from Urbana and Monmouth. The vertical progress of *Septoria* brown spot remained low for most of the season and didn’t reach halfway up the plant until 95 to 113 days after planting (approximately R5 to R6). Then, it reached 100% for most of the treatments at the last rating at R7 (Fig 1). The percent of chlorotic area was below 20% for all treatments until 114 days after planting (dap) (R6) but then increased to above 50% for the last rating at 125 dap (R7; S1 Fig). The necrotic area remained below 20% throughout the study. The defoliation rate increased gradually, starting at R1, and reached close to 100% by the last rating at R7. Due to the differences in disease progression between the traits, the final rating for vertical progress was considered to be R6, while for the other disease components were R7. AUDPC for vertical progress was calculated with the ratings from V7 to R6; for chlorotic and necrotic area with the ratings from V7 to R7; and for defoliation rate with the ratings from R1 to R7.

**Fig 1.**
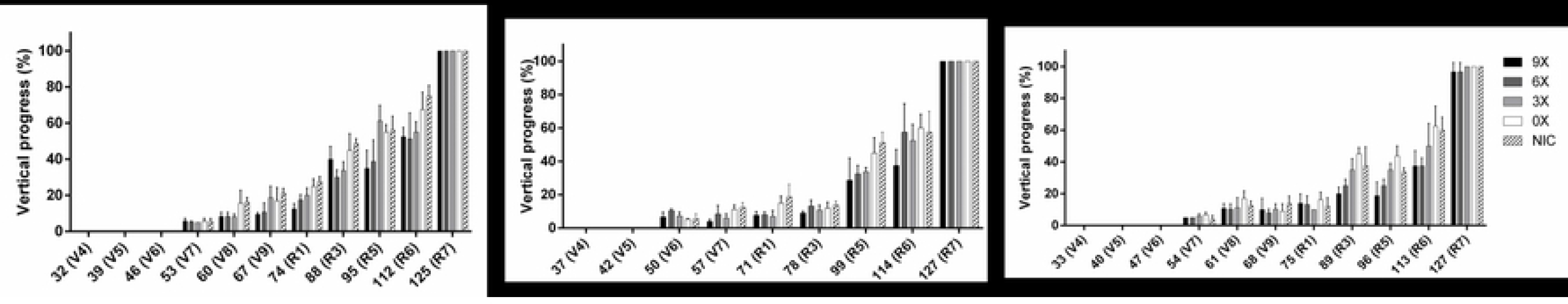
Vertical disease progress of *Septoria* brown spot from 32 to 127 days after planting on inoculated field trials in three locations in Illinois. Bars are the standard error.

Fungicide treatments significantly reduced the AUDPC for all four components of disease severity (*p* < 0.02) (S1 Table). At the end of the season, fungicide treatments significantly reduced the vertical progress at R6 (*p* = 0.008), chlorotic area at R7 (*p* = 0.004) (Fig 2) and showed a trend for necrotic area at R7 (*p* = 0.115) (S1 Table). Three, six, and nine weekly fungicide applications reduced 17% to 33% of the disease vertical progress as compared to the plots that were not treated with fungicide. For chlorotic area only six and nine weekly fungicide applications significantly reduced 18% and 21% of the chlorotic area as compared to the non-treated controls. No differences were observed between the inoculated and not-inoculated plots (Fig 2).

**Fig 2.**
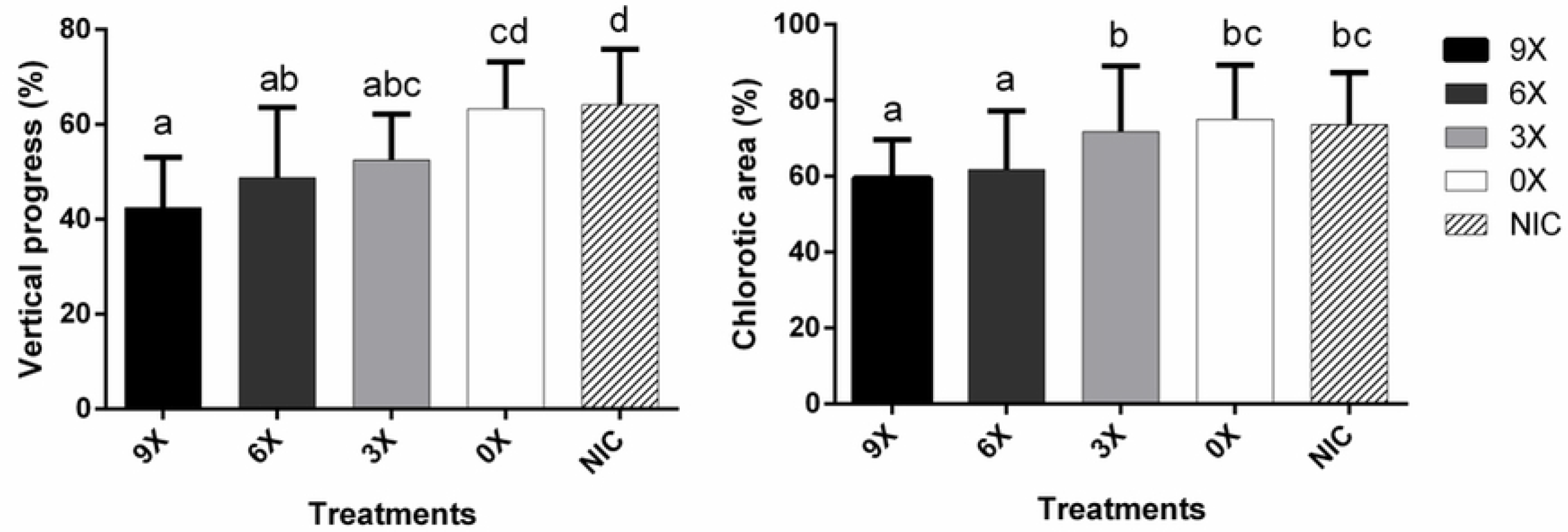
Fungicide treatments significantly reduced the vertical progress and chlorotic area caused by *Septoria* brown spot in three locations in Illinois. Bars are standard error, and letters above the bars are pairwise comparisons based on Fisher’s Least Significant Difference (LSD) method (*p* < 0.05).

For the disease traits, the largest variance was due to the location effects (Table 1), which ranged between 15% to 78%. Only a maximum of 3% of the variance was due to the block within location, and a maximum of 35% due to the location by treatment interaction. The one exception was for chlorotic area which had no variance due to location and 52% variance due to location by treatment interaction (Table 1).

**Table 1.** Variance components for *Septoria* brown spot response to fungicide application (two treatments: not inoculated control and three applications of chlorothalonil) for disease and yield components evaluated in three locations in Illinois.

| Disease or yield component | Location | Block<br>(location) | Location x<br>treatment | Residual |
| --- | --- | --- | --- | --- |
| Vertical progress (AUDPC) | 78% | 3% | 0% | 18% |
| Vertical progress (R6) | 15% | 2% | 5% | 78% |
| Chlorotic area (AUDPC) | 0% | 0% | 52% | 48% |
| Chlorotic area (R7) | 23% | 0% | 5% | 72% |
| Necrotic area (AUDPC) | 33% | 0% | 35% | 31% |
| Defoliation rate (AUDPC) | 75% | 0% | 10% | 15% |
| Yield | 79% | 0% | 6% | 16% |
| 100-seed weight | 75% | 8% | 8% | 9% |
| Pods per plant | 27% | 17% | 5% | 51% |
| Seeds per plant | 40% | 17% | 0% | 43% |

### 3.2 Yield response to fungicide treatments

Fungicide treatment did not have a significant effect on yield, yield response, 100-seed weight, pods per plant or seeds per plant. For yield and 100-seed weight, the largest percentage of variance was due to location effects (79% and 75% respectively). However, most of the variance was in the residual for pods per plant (51%) and seeds per plant (43%) (Table 1). A trend of higher yield on the plots treated nine times was observed in one of the locations (Perry). Two of the locations (Perry and Urbana) resulted in similar yields, but the yield was lower in Monmouth (Fig 3).

**Fig 3.**
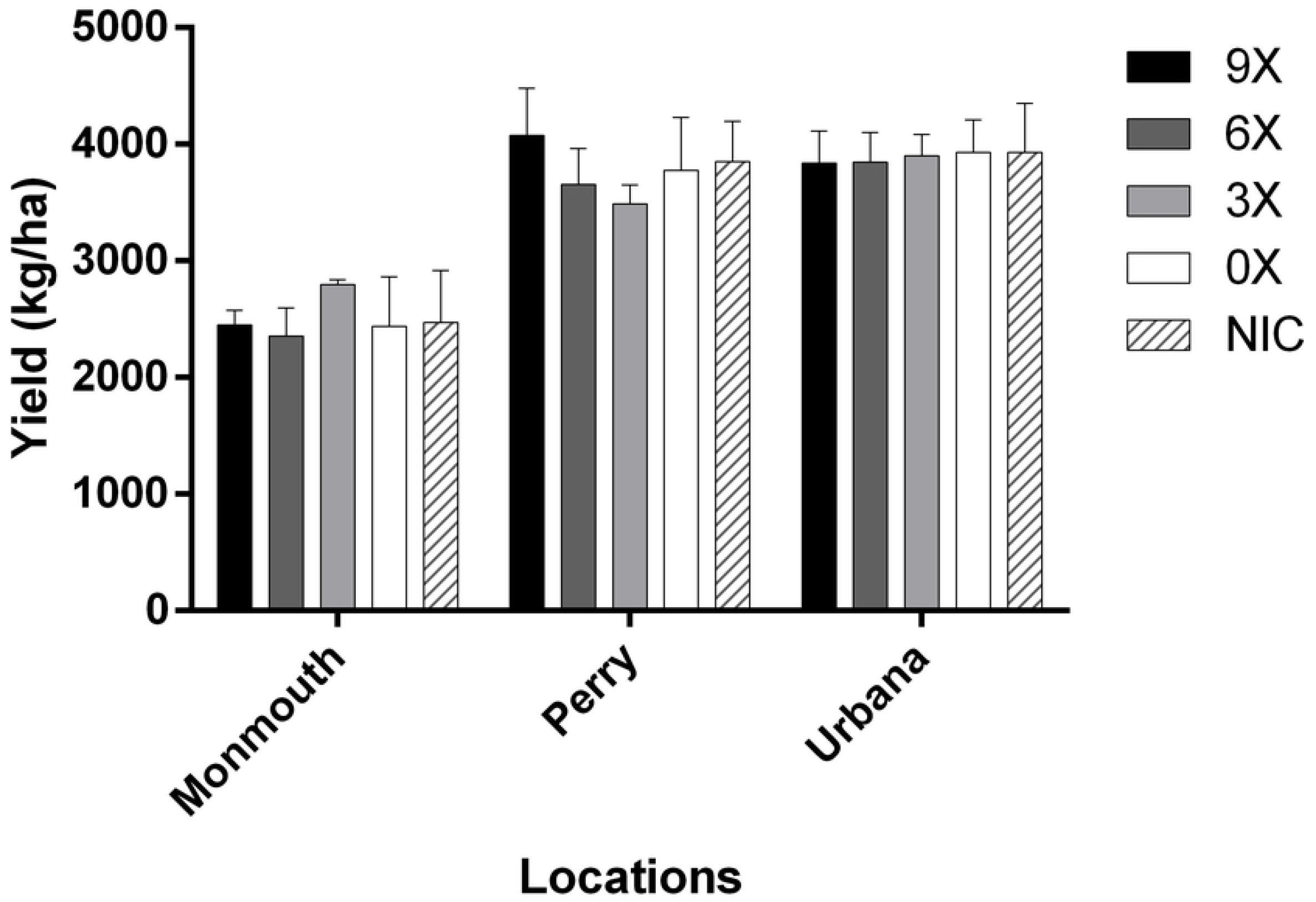
Mean yield (kg/ha) of field trials inoculated with *Septoria glycines* and treated with fungicide sprays for 9 times (9X), 6 times (6X), 3 times (3X), not sprayed (0X), and a not inoculated control (NIC) in three locations in Illinois. Bars are the standard error.

### 3.3 Correlation between brown spot disease and yield

Significant and week negative correlations were observed between yield and vertical progress (both AUDPC *r* = −0.36 and final rating at R6 *r* = −0.28), and a trend was observed between yield and the AUDPC of chlorotic area (*r* = −0.26; Table 2). Also, a negative correlation was observed between yield and 100-seed weight (*r* = −0.49). The AUDPC and the final rating of vertical progress (R6) were significantly correlated with the AUDPC of chlorotic area but not the final rating of chlorotic area (Table 2). The 100-seed weight in addition to being negatively correlated with yield, it was also negatively correlated with chlorotic area at R7 but positively correlated with the AUDPC of vertical progress. No significant correlations were found between yield response and yield, vertical progress, chlorotic area or 100-seed weight. No correlations were significant when the analysis was done by location (not shown).

**Table 2.** Pearson correlation coefficient (above) and their *p*-value (below) for selected components of disease and yield for the combined data of three locations.

|  |  | Vertical progress |  | Chlorotic area |  | 100-SW<sup>a</sup> | Yield response |
| --- | --- | --- | --- | --- | --- | --- | --- |
|  |  | AUDPC | R6 | AUDPC | R7 |  |  |
| Yield |  | -0.36 | -0.28 | -0.26 | 0.08 | -0.49 | 0.19 |
|  |  | 0.008 | 0.034 | 0.063 | 0.546 | <0.001 | 0.220 |
| Vertical Progress | AUDPC | - | 0.85 | 0.72 | 0.11 | 0.34 | -0.03 |
|  |  | - | <.001 | <.001 | 0.430 | 0.009 | 0.871 |
|  | R6 | - | - | 0.62 | 0.21 | 0.12 | -0.05 |
|  |  | - | - | <.001 | 0.128 | 0.371 | 0.768 |
| Chlorotic Area | AUDPC | - | - | - | 0.66 | 0.03 | -0.10 |
|  |  | - | - | - | <.001 | 0.804 | 0.561 |
|  | R7 | - | - | - | - | -0.38 | 0.03 |
|  |  | - | - | - | - | 0.005 | 0.838 |
| 100-SW<sup>a</sup> |  | - | - | - | - | - | 0.03 |
|  |  | - | - | - | - | - | 0.859 |

<sup>a</sup> Weight of 100 seeds

### 3.4 Linear regression analysis

Stepwise regression analysis identified the final rating and AUDPC of vertical progress to be significant predictor variables when included in the model independently. The estimated regression slope was −14.21 (*p* = 0.0357) for vertical progress rating data at R6, and −0.69 (*p* = 0.0007) for AUDPC of vertical progress. The coefficient of determination (R^2^) was 0.08 and 0.20, respectively (Fig 4). This indicated that 8% to 20% variation of yield was explained by the vertical progress of *Septoria* brown spot.

**Fig 4.**
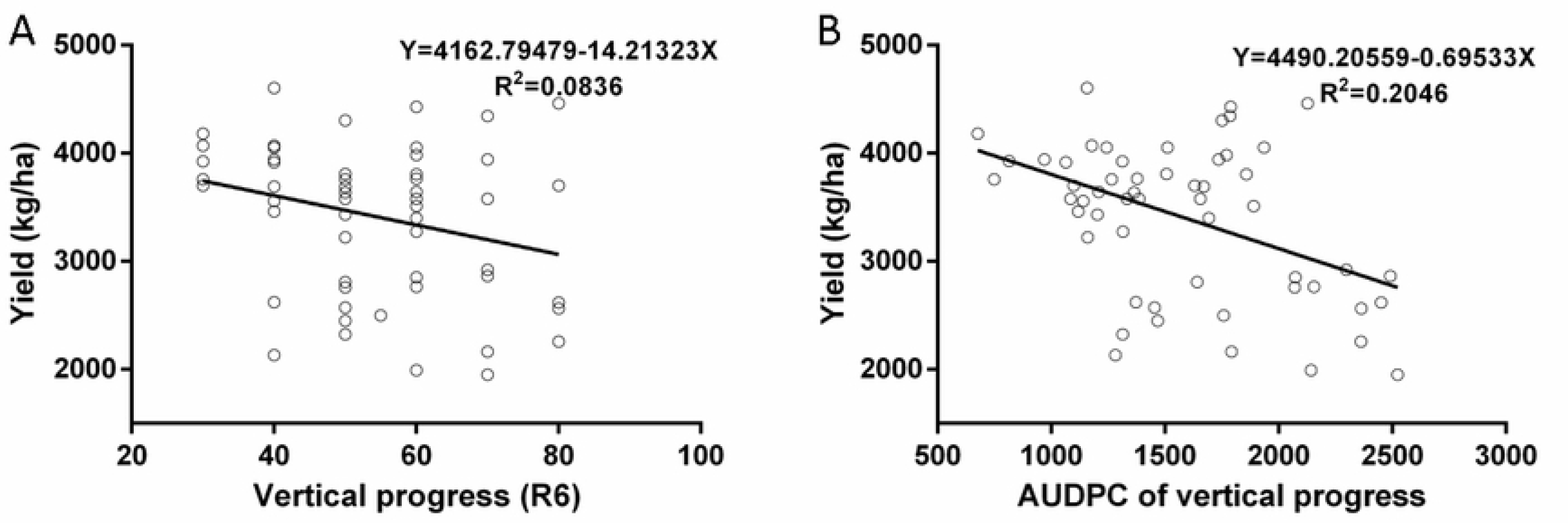
Linear relationship between yield and *Septoria* brown spot disease intensity (damage curve). (A) Vertical progress at R6. (B) AUDPC (V7 to R6) of vertical progress.

### 3.5 Power Analysis

Simulations based on our empirical data show that the number of locations required to obtain more than 80% statistical power to detect a yield reduction of 269 kg/ha (4 bsh/acr) between the untreated control and plots sprayed three times is 15 (Fig 5). Increasing the number of replicates within location did not result in any reasonable power increases (not shown). To obtain 80% power in a 3-location experiment, the difference between the untreated control and the plots sprayed three times would have to be 1008.75 kg/ha (15 bsh/acr). The number of locations required for other yield components was 16 for pods per plant (effect of 4 pods/plant), 6 for seeds per plant (effect of 9 seeds/plant), and 8 for 100-seed weight (effect of 0.6g/100-seeds) (not shown).

**Fig 5.**
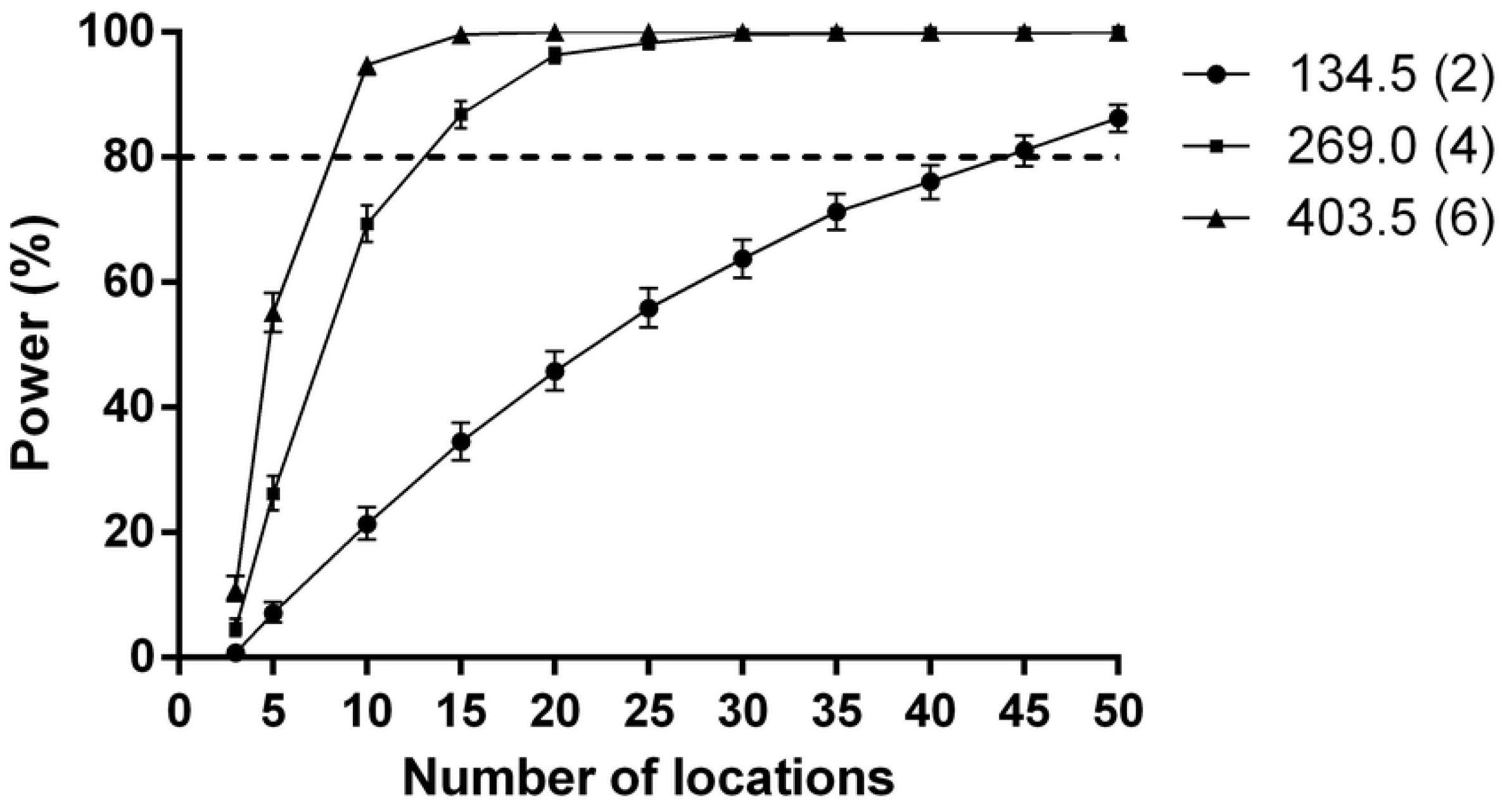
Power curves simulated based on our data for the required number of locations to detect statistically significant yield response (80% power) for treatment differences (effect) of 134.5 (2), 269.0 (4), and 403.5 (6) kg/ha (bsh/acr) on inoculated experiments.

## 4 Discussion

Many foliar diseases affect soybean through the growing season, and in many cases, foliar fungicide applications might be useful to protect yields from these diseases. One of the most common foliar diseases of soybean is *Septoria* Brown Spot. Despite it being widely distributed, the effects of *Septoria* brown spot on yield remain poorly characterized due to the variation of the results within studies [4]. Better recommendations for fungicide application due to any disease can be obtained with precise understanding of the yield limiting effect of each pathogen. It has been long know that under low disease pressure, the estimation of the disease effect on yield has high standard errors on the mean estimates [25]. Thus, to start untangling the effects of late-season soybean disease on yield, we tackled the most prevalent of the diseases: *Septoria* brown spot. We conducted multi-location inoculated field trials to generate empirical data of disease effects and model the linear relationship of the damage due to this disease.

In this study, different levels of disease severity of brown spot were obtained at three locations following three, six, or nine weekly applications of chlorothalonil after disease inoculation. Among the four disease components investigated (vertical progress, necrotic area, chlorotic area, and defoliation rate), vertical progress was the best trait to identify differences in the development of brown spot during the reproductive stages. Conversely, necrotic area remained low throughout the season while chlorotic area and defoliation rate did not show any variation until R7 or later.

Diseased leaf area (DLA) is one of the standard methods used to evaluate brown spot severity [10, 11, 26]. Pataky and Lim (1981) [9] used DLA, vertical progress and defoliation rate to evaluate the disease severity in a 2-year experiment. In inoculated plots, DLA reached 50-60% at R6 stage; vertical progress reached 50% at R6, and over 90% at the last rating in both years; the defoliation rate was about 30% at R6, and reached 60% at the last rating only in one year. Kamicker and Lim (1985) [14] also evaluated DLA, vertical progress and defoliation rate at R6. DLA was between 0-20% while the vertical progress and defoliation rate was presented by node, not percentage, which makes it difficult to compare with other studies. Cruz et al. (2010) [12] used image analysis software to calculate the DLA from lower and middle canopy leaves. The DLA was below 20% in both lower and middle canopies at the reproductive stage. In our study, the vertical progress reached about 60% in the control plots (no fungicide applied), which is similar to the results from Pataky and Lim (1981) [9]. The necrotic area and chlorotic area in our results were relatively lower than the results from Pataky and Lim (1981) [9] but similar to Cruz et al. (2010) [12] and Kamicker and Lim (1985) [14]. As for the defoliation rate, the trend is similar to the results in Pataky and Lim (1981) [9]. Both vertical progress and DLA appear to be adequate to evaluate this disease and enable comparisons to previous studies. Our study shows that the multiple application of fungicide resulted on different levels of disease severity. However, there were no significant differences between the inoculated (0X) and not inoculated control (NIC) plots. This might indicate the inoculation method failed, or that there were high levels of natural inoculum in the field.

Although different levels of disease severity were obtained, no significant difference in yield was found between treatments in this study. Both Lim (1980) [11] and Cooper (1989) [16] obtained consistent yield response to fungicide application. Lim (1980) [11] reported a 12-34% yield increase between fungicide-treated and inoculated plots and 8% of yield increase between fungicide-treated and naturally infected plots in a 2-year experiment. Cooper (1989) [16] reported 2.2% to 15.4% yield increase in the fungicide-treated plots in a 3-year experiment. However, other studies only obtained yield response in one year or even no yield response throughout the experiment. Young and Ross (1978) [26] reported that 13.4 to 17.8% (404-514 kg/ha) yield reduction was found in only one year (out of 2 tested) between control and inoculated plots. Pataky and Lim (1981) [9] reported 16% (30 mg) reduction of 100-seed weight in the inoculated plot only in one year (out of 2 tested). Kamicker and Lim (1985) [14] reported no yield reduction found in inoculated plots in a 2-year experiment. Cruz et al. (2010) [12] reported that the yield increased with the application of fungicides between 183.7 to 490 kg/ha (~4-10% increase) for 3 out of 6 location/years. The inconsistent results among the studies might be due to low disease severity [27], late appearance of the disease [14, 28] or insufficient statistic power for the experiment design [23].

The lack of significant effects for yield in our study prompted us to conduct a more detailed analysis of the sources of variation and a power analysis. Statistic power is the probability to detect a significant treatment effect in a study. Small-plots trials are commonly used since the small-plots are easier to handle (e.g., disease evaluation and sample collection) and lower cost. Kandel et al. (2018) [23] compared the yield response between small-plots trials and on-farm replicated strip trials and found that the trial type was not significantly different. However, Kandel et al. (2018) [23] point out the importance of the statistical power in small-plots trials when the disease pressure is low. The results from our power analysis indicate that we needed between eight and 13 environments to detect 269 to 807 kg/ha (4 to 6 bsh/acr; 7-12%) yield reduction effects due to *Septoria* brown spot. Our results agree with the result In Kandel et al. (2018) [23] that the significant yield difference below 134-201 kg/ha (2-3 bu/ac) in small-plot trials was difficult to detect.

A significant negative correlation was found between yield and vertical progress of the disease. Eight to 20% of the variation in yield was explained by the rating data at R6 or the AUDPC of vertical progress. From the linear model, when the vertical progress of brown spot at R6 increase by 10%, the yield decreases by 142.13 kg/ha (2 bushels/acre; 3.4%). Cruz et al. (2010) [12] also reported that the disease severity at R5 or R6 explains 3 to 10% variation of the yield response. In Young et al. (1978) [26], the negative correlation between yield and disease severity was found in one year. Lim (1980) [11] reported a significant correlation between yield reduction and disease severity or AUDPC at R6 stage. This shows that brown spot of soybean plays a role in limiting the yield, but the yield response is highly related to the location which vary due to weather conditions. In future experiments, we will need to include more locations to fine-tune the model for yield limitations of *Septoria* brown spot and include other late-season diseases. This information will help to define accurate thresholds for the timing of fungicide application in Illinois.

## Acknowledgments

We are grateful to Juan Pablo Granda Cruz for technical assistance.

## Supporting information

**S1 Table. Analysis of variance for the effect of fungicide treatments on multiple components of disease severity of *Septoria* brown spot at the end of the season and on their AUDPC values.**

**S1 Fig. Components of disease of *Septoria* brown spot of soybean from 32 to 127 days after planting on inoculated field trials in three locations in Illinois. Bars are the standard error.**

## References

1. USDA. World agricultural production. 2019; 1:32. Available from: https://apps.fas.usda.gov/psdonline/circulars/production.pdf

2. Hartman GL, Rupe JC, Sikora EJ, Domier LL, Davis JA, Steffey KL. Compendium of soybean diseases and pests, 5th ed. St. Paul, Minnesota, USA: The American Phytopathological Society; 2015.

3. Allen TW, Bradley CA, Sisson AJ, Byamukama E, Chilvers MI, Coker CM, et al. Soybean yield loss estimates due to diseases in the United States and Ontario, Canada, from 2010 to 2014. Plant Health Prog. 2017;18(1):19–27.

4. Carmona M, Sautua F, Perelman S, Reis EM, Gally M. Relationship between late soybean diseases complex and rain in determining grain yield responses to fungicide applications. J. Phytopathol. 2011;159(10):687–693.

5. Wolf FA. Report of the division of plant pathology. North Carolina Agricultural Experiment Station Annual Report. 1923;46:92.

6. Fehr WR, Caviness CE, Burmood DT, Pennington JS. Stage of development descriptions for soybeans, *Glycine max* (L.) Merrill. Crop Sci. 1971;11(6):929–931.

7. Mueller D, Wise K, Sisson A, Smith D, Sikora E, Bradley C, et al. A farmer’s guide to soybean diseases. St. Paul, Minnesota, USA: American Phytopathological Society Press; 2016.

8. Hemmi T. A new brown-spot disease of the leaf of *Glycine hispida* Maxim. caused by *Septoria glycines* sp.n. Trans. Sapporo Nat. Hist. Soc. 1915;6:12–17.

9. Pataky LK, Lim SM. Effect of *Septoria* brown spot on the yield components of soybeans Plant Dis. 1981;65:588–590.

10. Lim SM. Evaluation of soybean for resistance to *Septoria* brown spot. Plant Dis Rep. 1979;63:242–245.

11. Lim SM. Brown spot severity and yield reduction in soybean. Phytopathology. 1980;70(10):974–977.

12. Cruz CD, Mills D, Paul PA, Dorrance AE. Impact of brown spot caused by *Septoria glycines* on soybean in Ohio. Plant Dis. 2010;94(7):820–826.

13. Young LD, Ross JP. Resistance evaluation and inheritance of a nonchlorotic response to brown spot of soybean. Crop Sci. 1978;18:1075–1077.

14. Kamicker TA, Lim SM. Field evaluation of pathogenic variability in isolates of *Septoria glycines*. Plant Dis. 1985;69:744–746.

15. Pataky JK, Lim SM. Efficacy of benomyl controlling *Septoria* brown spot of soybean. Phytopathology. 1980;71:438–442.

16. Cooper RL. Soybean yield response to benomyl fungicide application under maximun yield conditions. Agronomy J. 1989;81:847–849.

17. Van Roekel RJ, Purcell LC. Understanding and increasing soybean yields. Proceedings of the Integrated Crop Management Conference. 2016;4. Available from: https://lib.dr.iastate.edu/cgi/viewcontent.cgi?referer=https://www.google.com/&httpsredir=1&article=1213&context=icm

18. Felipe de Mendiburu. CRAN—Package agricolae. 2017. [cited 12 Apri 2019]. Available from: https://cran.r-project.org/web/packages/agricolae/index.html

19. R Core Team. R: A Language and Environment for Statistical Computing. 2011 [cited 12 Apri 2019]. Available from: https://cran.r-project.org/bin/windows/base/

20. SAS Institute. The SAS system for Windows. Release 9.4. SAS Inst., Cary, NC. 2013 [cited 12 Apri 2019]. Available from: https://www.sas.com/en_us/software/sas9.html

21. GraphPad Software Inc. GraphPad Prism version 6 for Windows., La Jolla California., USA. 2016 [cited 12 Apri 2019]. Available from: https://www.graphpad.com/

22. Green P, MacLeod CJ, Nakagawa S. SIMR: an R package for power analysis of generalized linear mixed models by simulation. Methods Ecol Evol. 2016;7(4):493–498.

23. Kandel YR, Hunt CL, Kyveryga PM, Mueller TA, Mueller DS. Differences in small plot and on-farm trials for yield response to foliar fungicide in soybean. Plant Dis. 2018;102(1):140–145.

24. Kandel YR, Mueller DS, Hart CE, Bestor NRC, Bradley CA, Ames KA, et al. Analyses of yield and economic response from foliar fungicide and insecticide applications to soybean in the north central United States. Plant Health Prog. 2016.

25. James WC. Assessment of plant diseases and losses. Annu Rev Phytopathol. 1974;12(1):27–48.

26. Young LD, Ross JP. Brown spot development and yield response of soybean inoculated with *Septoria glycines* at various growth stages. Phytopathology. 1978;68(1):8–11.

27. Hershman DE, Vincelli P, A. KC. Foliar fungicide use in corn and soybeans. University of Kentucky Plant Pathology Fact Sheet. 2011; 1:9. Available from: https://plantpathology.ca.uky.edu/files/ppfs-gen-12.pdf

28. Henry RS, Johnson WG, Wise KA. The impact of a fungicide and an insecticide on soybean growth, yield, and profitability. Crop Prot. 2011;30(12):1629–1634.

